# A neural network for religious fundamentalism derived from patients with brain lesions

**DOI:** 10.1101/2023.12.22.572527

**Authors:** Michael A. Ferguson, Erik W. Asp, Isaiah Kletenik, Daniel Tranel, Aaron D. Boes, Jenae M. Nelson, Frederic L.W.V.J. Schaper, Shan Siddiqi, J. Seth Anderson, Jared Nielsen, James R. Bateman, Jordan Grafman, Michael D. Fox

## Abstract

Religious fundamentalism, characterized by rigid adherence to a set of beliefs putatively revealing inerrant truths, is ubiquitous across cultures and has a global impact on society. Understanding the psychological and neurobiological processes producing religious fundamentalism may inform a variety of scientific, sociological, and cultural questions. Research indicates that brain damage can alter religious fundamentalism. However, the precise brain regions involved with these changes remain unknown. Here, we analyzed brain lesions associated with varying levels of religious fundamentalism in two large datasets from independent laboratories. Lesions associated with greater fundamentalism were connected to a specific brain network with nodes in the right orbitofrontal, dorsolateral prefrontal, and inferior parietal lobes. This fundamentalism network was strongly right hemisphere lateralized and highly reproducible across the independent datasets (*r* = 0.82) with cross-validations between datasets. To explore the relationship of this network to lesions previously studied by our group, we tested for similarities to twenty-one lesion-induced conditions. Lesions associated with confabulation and criminal behavior showed a similar connectivity pattern as lesions associated with greater fundamentalism. Moreover, lesions associated with poststroke pain showed a similar connectivity pattern as lesions associated with lower fundamentalism. These findings are consistent with hemispheric specializations in reasoning and lend insight into previously observed epidemiological associations with fundamentalism, such as cognitive rigidity and outgroup hostility.

## Introduction

The basis of one’s religious beliefs and behaviors has been an enduring focus of psychological inquiry and, more recently, neuroscientific investigation^1,2^. One facet of religiosity is religious fundamentalism: an adherence to religious doctrines believed to be inerrant, a devotion to religious practices considered immutable, and a perceived special relationship with a deity ^3,4^. Fundamentalism is associated with authoritarianism^5,6^, cognitive rigidity^7–9^, lower complexity of thought on religious issues^10,11^, a reduced likelihood of doubt^6,11,12^ increased acceptance of misinformation^13,14^, delusion-like ideation^15^, anti-intellectualism^15,16^, prejudicial attitudes^3,17^, and outgroup hostility^18^. Fundamentalism may confer group-level advantages: including stronger ingroup commitment^19^ and an increased sense of belonging and well-being^20^.

Research examining the determinants of religious fundamentalism has often focused on environmental or socialization factors^21^ such as socioeconomic and family affiliation variables^22,23^. However, neuroscientific and behavioral genetic research argue for a biological influence toward religious experiences and attitudes: pharmacological, neuroimaging, and psychophysiological studies indicate that unique patterns of neural activity are correlated with religious states^24–28^ and twin studies show religious fundamentalism to be strongly heritable^29–31^. Thus, neurobiological factors likely predict a style of cognitive and emotional processing that tends to result in fundamentalist attitudes^12,13^. The cognitive and behavioral associations with fundamentalism may be clarified by the identification of brain regions or networks underlying religious fundamentalism^32^.

Brain lesion studies allow for causal inferences into human behavior^33,34^, including religiosity^12,2,35^. Prior research has shown that damage to the prefrontal cortex is associated with higher levels of religious fundamentalism^6,7^. However, this preliminary work was limited by sample size and a narrow focus on the prefrontal cortex. The prefrontal cortex is a large and highly connected region^36^ that participates in a number of brain-wide networks^36,37^. Neurobiological factors that underlie a cognitive or emotional style of processing associated with religious fundamentalism are most likely a distributed brain network phenomenon^34^. To date, no studies have investigated fundamentalism using a brain network approach. As a result, there remains a critical gap in the evidence relating to religious fundamentalism and the brain.

Here, we examined two large and independently-collected human lesion datasets of rare neurological patients (*N*1 = 106; *N*2 = 84) to determine whether religious fundamentalism could be mapped to a functional network across the brain. We used an analytic method, lesion network mapping, which relies on functional connectivity data from healthy individuals to infer functional networks that may be disrupted by a lesion^2,38^. This technique allowed an examination of our results with our extensive database of lesions associated with a variety of behavioral, neurological, and psychiatric conditions (*N*3 = 899).

## Results

Lesions associated with religious fundamentalism occurred in multiple different brain regions across our two datasets (Figure 1) emphasizing the utility of our brain network approach. We combined both independent datasets and performed voxelwise statistical tests for lesion network associations with Religious Fundamentalism Scale^2^ (RFS) scores (*N*_combined_ = 190). Functional connectivity between each lesion location and the rest of the brain was computed using a publicly available normative connectome dataset from 1000 healthy right-handed subjects (Figure 2). Connections significantly associated with RFS scores were identified (Figure 3, FWE p_corrected_ < 0.05). Higher fundamentalism scores were associated with positive connectivity between lesion locations and right superior orbital frontal, right middle frontal, right inferior parietal, right inferior temporal cortices, and left cerebellum (Figure 3b; see Table 1). The fundamentalism network maxima clusters were strongly right lateralized in the cerebral cortex. We also observed left cerebellum maxima clusters resulting from contralateral cerebro-cerebellar connections^39^. Lower fundamentalism scores were associated with negative connectivity between lesion locations and the left paracentral lobule, right cerebellum, and bilateral middle temporal cortices (Figure 3c; see Table 1).

**Figure 1:**
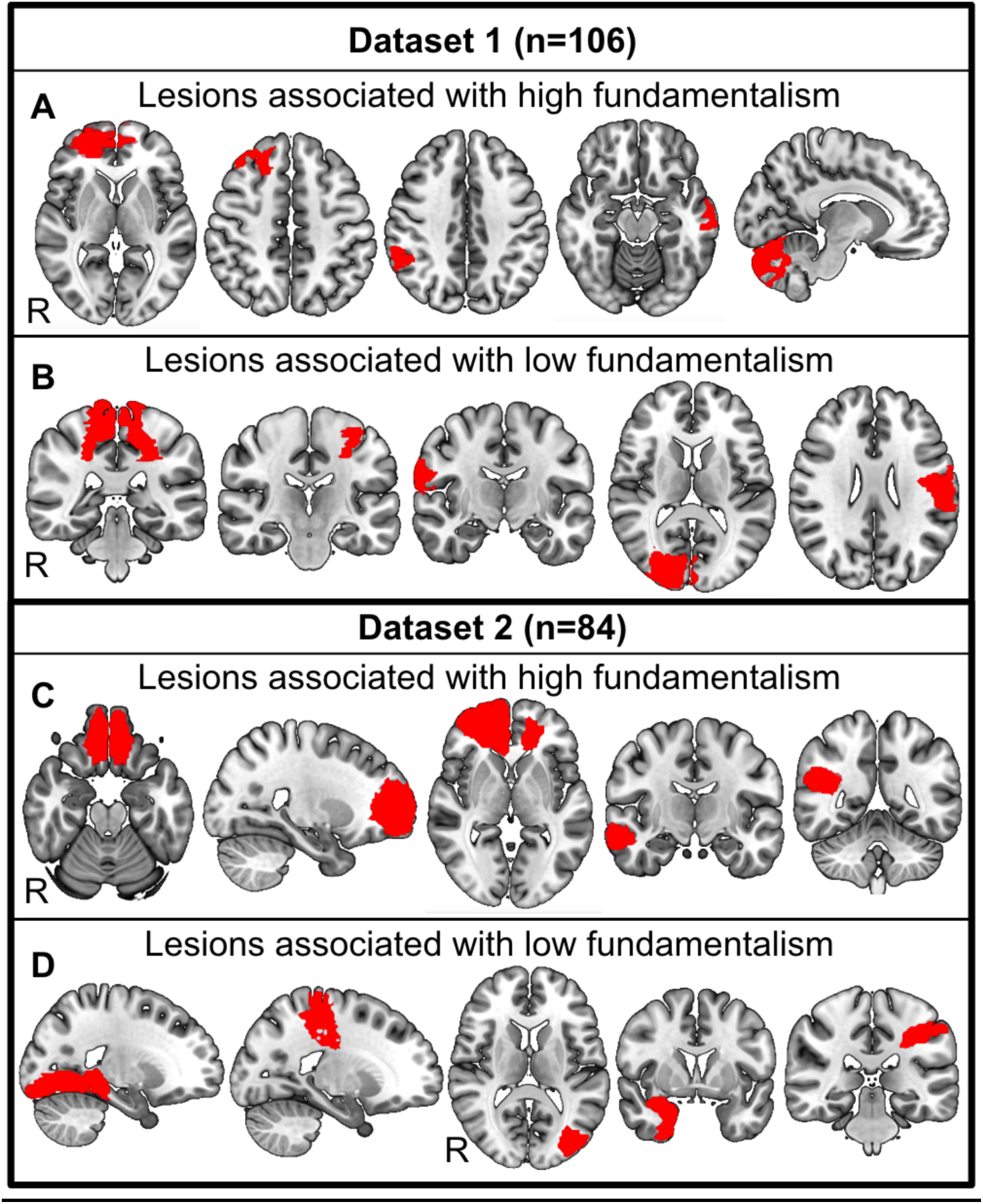
Lesions associated with religious fundamentalism occur in many different brain regions beyond the prefrontal cortex. (A) Brain lesions of five patients with the highest self-report scores for religious fundamentalism from dataset 1. (B) Brain lesions of five patients with the lowest self-report scores for religious fundamentalism from dataset 1. (C) Brain lesions of five patients with the highest self-report scores for religious fundamentalism from dataset 2 (D) Brain lesions of five patients with the lowest self-report scores for religious fundamentalism from dataset 2. Note that lesions associated with high and low fundamentalism are highly heterogenous across brain regions and datasets.

**Figure 2:**
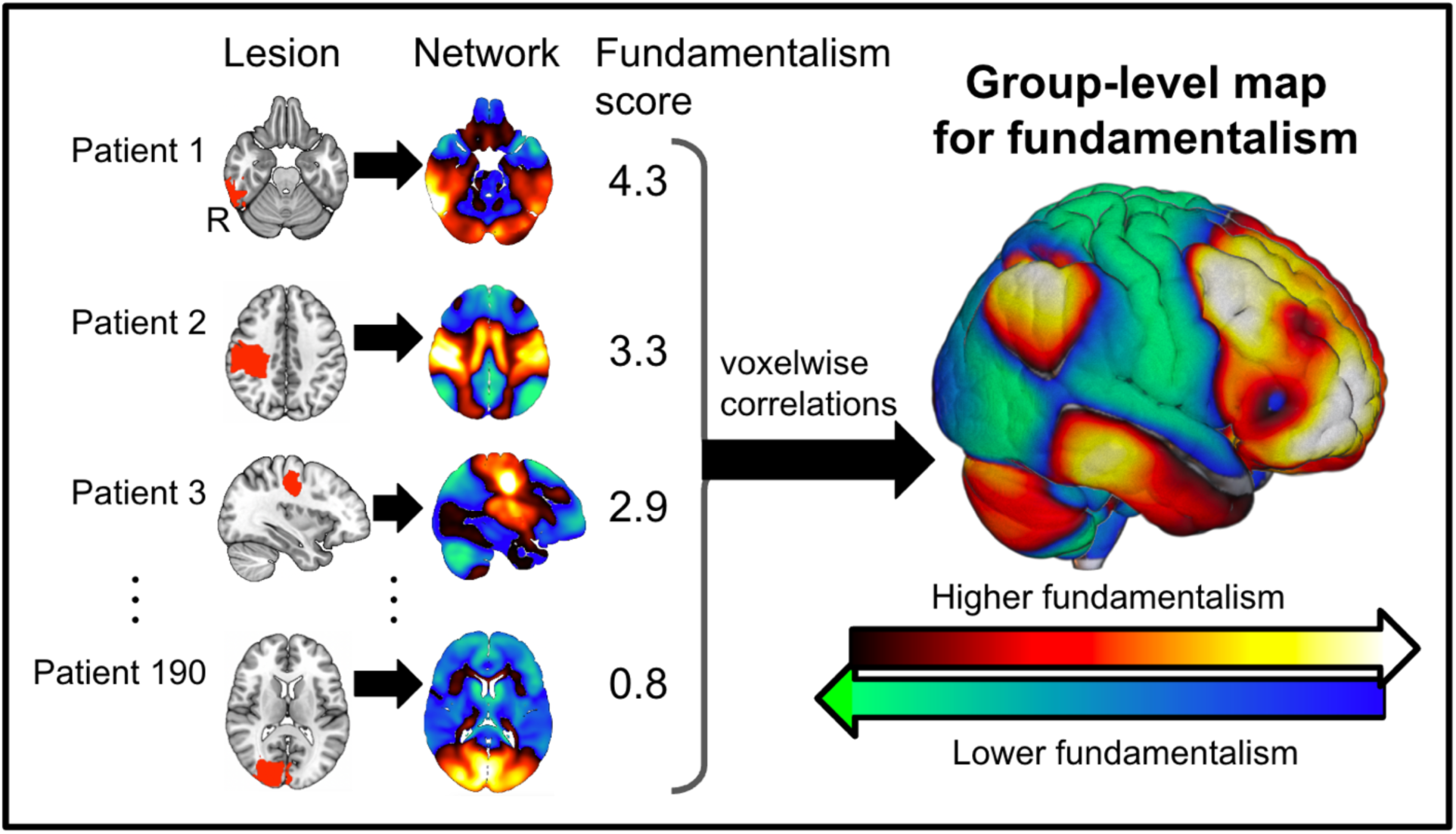
Data-driven method for identifying a lesion network for religious fundamentalism. The network of brain regions functionally connected to each lesion was computed for dataset 1 using resting-state functional connectivity data from a large database of healthy volunteers (*N* = 1000). Lesions and lesion networks are shown for 4 of the 190 patients in the combined datasets (*N*_combined_ = 190). Positively connected voxels are shown in warm colors, while negatively connected voxels are shown in cool colors. Connections associated with religious fundamentalism scores were then identified (right). Warm colors in the group-level map indicate that functional connectivity with lesions is more likely associated with higher scores for religious fundamentalism. Conversely, cool colors in the group-level map indicate that functional connectivity with brain lesions is more likely associated with lower scores for religious fundamentalism.

**Figure 3:**
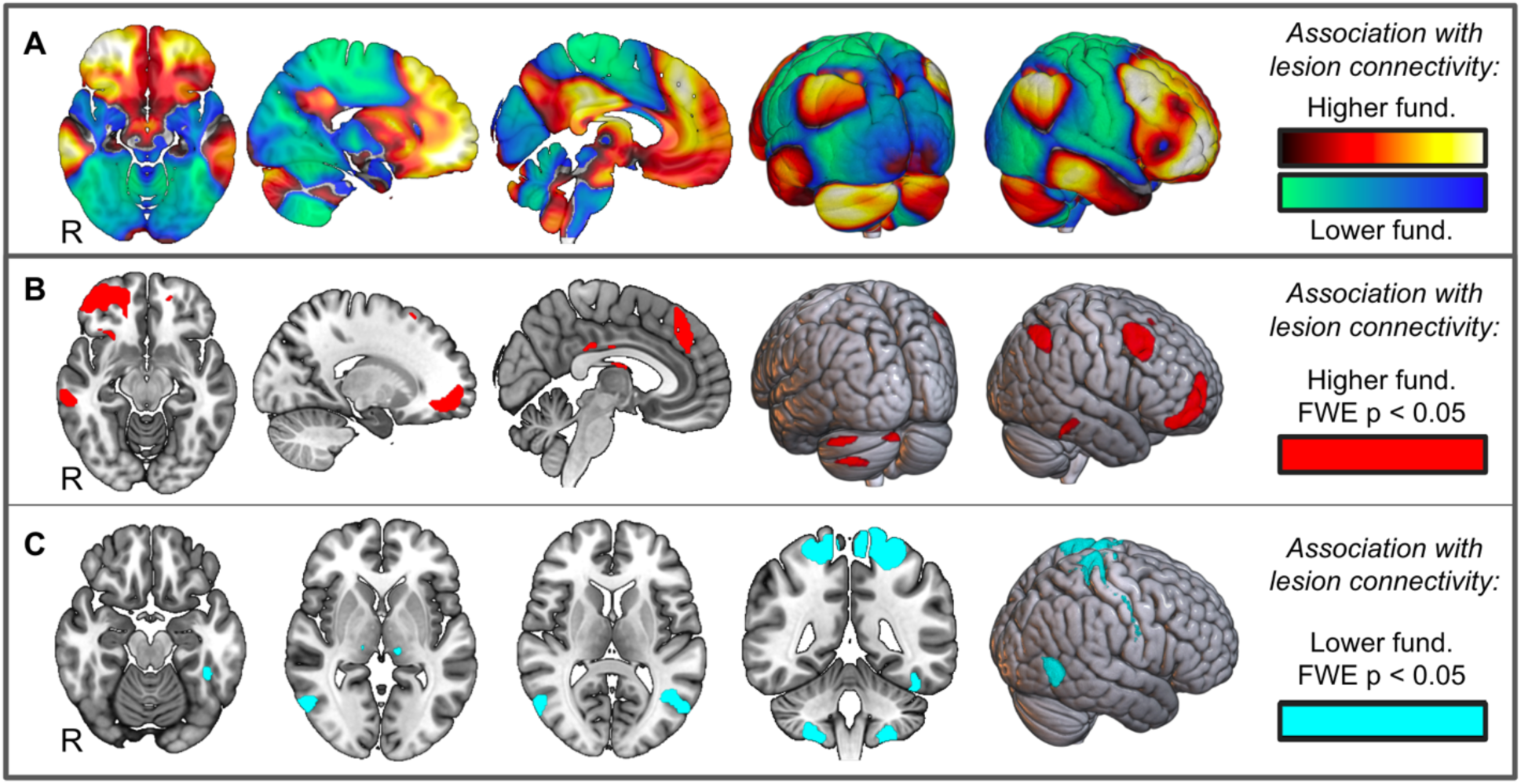
Religious fundamentalism network derived from combined datasets (*N* = 190). (A) Whole brain topography (unthresholded) for statistical associations between brain lesion connectivity and religious fundamentalism scores. Warm colored regions indicate positive functional connectivity relative to lesions associated with higher religious fundamentalism. Cool colored regions indicate negative functional connectivity relative to lesions associated with higher religious fundamentalism. (B) Voxels with statistically significant connectivity to brain lesions associated with higher religious fundamentalism following family-wise error multiple comparison correction (FWE p < 0.05). (C) Voxels with statistically significant connectivity to brain lesions associated with lower religious fundamentalism following family-wise error multiple comparison correction (FWE p < 0.05).

**Table 1:** Locations of local maxima and local minima within our lesion-derived brain network for religious fundamentalism. All clusters achieved significance at a family-wise error (FWE) multiple comparison correction of p < 0.05.

| Cluster size<br>(2 mm voxels) | Peak<br>*t* | MNI coordinates | Peak Structure |
| --- | --- | --- | --- |
| 13787 | 5.1 | [18, 50, -11] | R Superior Orbital Frontal |
| 6020 | 5 | [46, 14, 46] | R Middle Frontal |
| 3200 | 4.8 | [50, -58, 56] | R Inferior Parietal |
| 2414 | 4.8 | [4, 38, 40] | R Superior Medial Frontal |
| 2273 | 4.6 | [-35, -76, -45] | L Crus 2, Cerebellum |
| 2001 | 4.9 | [60, -28, -16] | R Inferior Temporal |
| 1458 | 4.9 | [-12, -86, -26] | L Crus 2, Cerebellum |
| 768 | 4.7 | [-46, -76, -28] | L Crus 1, Cerebellum |
| 507 | 4.5 | [-20, 52, -10] | L Superior Orbital Frontal |
| 298 | 4.5 | [2, -30, 30] | R Middle Cingulum |
| 265 | 4.5 | [14, 6, 20] | R Caudate |
| 192 | 4.5 | [2, -12, 29] | R Middle Cingulum |
| 147 | 4.4 | [20, 23, 60] | R Superior Frontal |
| 135 | 4.4 | [26, 22, -11] | R Insula |
| 95 | 4.4 | [4, -8, 16] | R Fornix |
| 33770 | -5.2 | [10, -36, 76] | L Paracentral Lobule |
| 3010 | -4.8 | [22, -50, -56] | R Cerebellum (8) |
| 2494 | -4.7 | [-54, -68, 12] | L Middle Temporal |
| 2162 | -4.8 | [56, -66, 4] | R Middle Temporal |
| 1671 | -4.8 | [-26, -42, -56] | L Cerebellum (8) |
| 1181 | -4.7 | [-42, -40, -24] | L Fusiform Gyrus |
| 409 | -4.4 | [30, -16, 62] | R Precentral Gyrus |
| 154 | -4.5 | [-12, -26, 0] | L Thalamus |
| 55 | -4.3 | [14, -19, 74] | R Precentral Gyrus |

Notably, religious fundamentalism whole brain association maps derived independently from dataset 1 and 2 robustly replicated each other, demonstrating a strong spatial correlation (*r =* 0.82, *p =* 0.02; Figure 4a, 4b). Cross-validation analyses further indicated the reliability of the fundamentalism network results between the independently-collected lesion datasets. Correlations were observed between RFS scores for lesions in dataset 1 and the similarity of dataset 1 patient lesions with the dataset 2 group-level fundamentalism network (*r =* 0.28, *p =* 0.003). Correlations were also observed between RFS scores for patients in dataset 2 and the similarity of dataset 2 patient lesions with the group-level fundamentalism network derived independently from dataset 1 (*r =* 0.27, *p =* 0.01; Figure 4c, 4d). A secondary cross-validation analysis quantified the amount of spatial overlap between each patient’s lesion relative to a group-level fundamentalism map derived from the opposing dataset. Correlations were observed between RFS scores for patients in dataset 1 and the spatial overlap of dataset 1 patient lesions to the dataset 2 group-level fundamentalism network (*r =* 0.21, *p =* 0.03). Correlations were also observed between RFS scores for patients in dataset 2 and the spatial overlap of dataset 2 patient lesions with the group-level fundamentalism network derived independently from dataset 1 (*r =* 0.29, *p =* 0.008; Figure 4e, 4f). Finally, to test the utility of our lesion network mapping approach we performed a data-driven analysis of lesion locations (rather than lesion networks) on our data. Voxel lesion-symptom mapping (VLSM) analyses failed to reveal significant results. The spatial correlation between the VLSM maps from the two datasets was not greater than would be predicted by chance (*r =* 0.069, *p =* 0.82; see Supplementary Methods and Supplementary Figure 1 for VLSM statistical approach and results).

**Figure 4:**
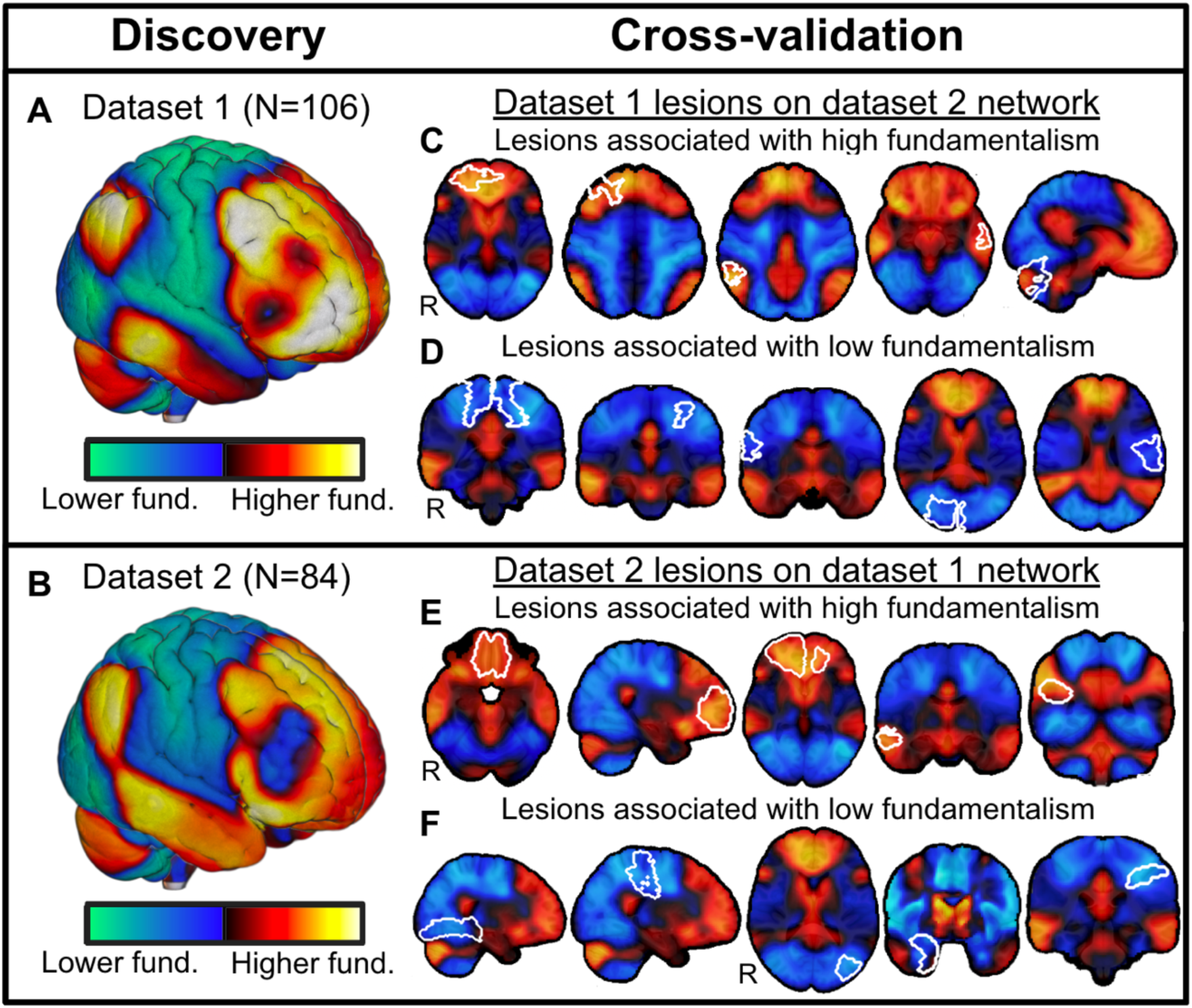
Cross-validation of lesion network mapping results across two independent datasets. (A) Discovery: lesion network mapping in dataset 1 defined a whole brain statistical map that relates lesion connectivity with religious fundamentalism scores. Positive correlations between lesion connectivity and religious fundamentalism scores are displayed in warm colors, and negative correlations between lesion connectivity and religious fundamentalism scores are displayed in cool colors. (B) Replication: lesion network mapping of religious fundamentalism in dataset 2 demonstrated a strong spatial similarly to lesion network map for religious fundamentalism from dataset 1 (spatial correlation, *r =* 0.82, *p =* 0.02). Cross-validations: (C) Brain lesions from dataset 1 (white outlines) associated with high religious fundamentalism scores intersect the lesion network connections from dataset 2 that were also associated with high religious fundamentalism scores (shown in warm colors). (D) Conversely, brain lesion from dataset 1 (white outlines) associated with low religious fundamentalism scores intersect the lesion network connections from dataset 2 that were also associated with low religious fundamentalism scores (shown in cool colors; *r =* 0.21, *p =* 0.03). (E) and (F) The same cross-validation analysis was also significant when brain lesion locations from dataset 2 (white outlines) were compared to whole brain map of lesion network correlations with religious fundamentalism derived from dataset 2 (*r =* 0.29, *p =* 0.008).

Religious fundamentalism in healthy individuals has a number of epidemiological associations including cognitive rigidity, reduced likelihood to doubt, prejudice, and outgroup hostility. If neurobiological factors influence a cognitive and emotional processing style that tends to result in fundamentalist attitudes, then these factors may also support these epidemiological associations. To investigate, we examined spatial similarities between our fundamentalism network and connectivity patterns for brain lesions associated with 21 behavioral, neurological, and psychiatric conditions (*N*3 = 899; see Figure 5 for complete condition list). Strong spatial similarities were observed between our religious fundamentalism network and brain lesion connectivity associated with pathological confabulation^40^ (spatial correlations: t_24_ = 8.3, *p* < 10^-7^, 95% CI, *r* = 0.32 to 0.53; Figure 5). Confabulation is the generation of conspicuously false beliefs or memories without the intent to deceive^41^ and is strongly associated with cognitive rigidity or perseveration during neuropsychological tasks^42,43^. Confabulation is also associated with failure to inhibit or doubt inaccurate memory elements during retrieval^12,41^. These aspects of confabulation are shared by healthy individuals high in fundamentalism, who also show cognitive rigidity and a reduced likelihood of doubt^6,8,9^.

**Figure 5:**
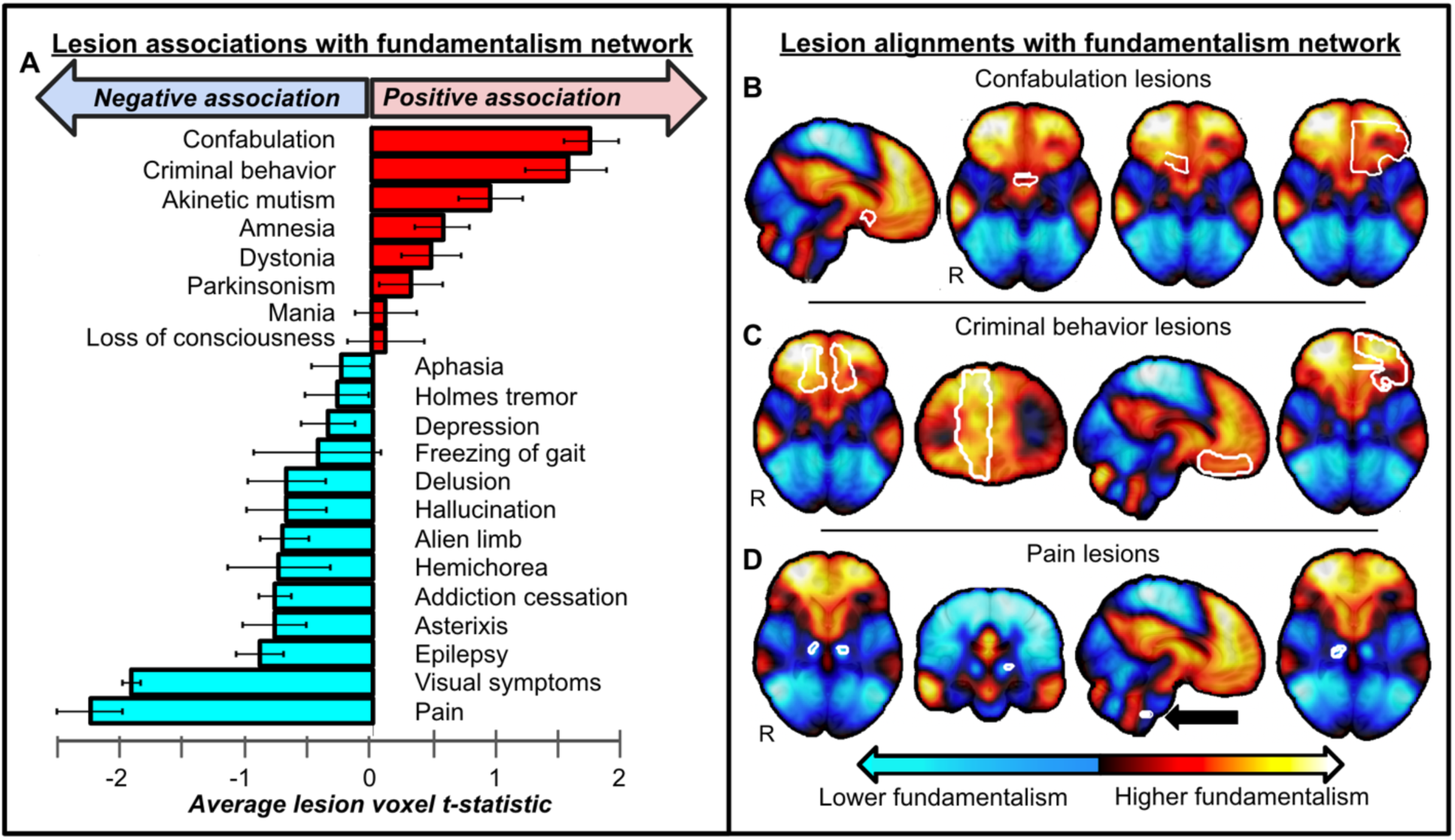
Lesions associated with behavioral, neurological, and psychiatric conditions intersect our religious fundamentalism circuit. (A) The average of voxel intensities within lesion locations associated with 21 different conditions (*N* = 899) are shown in a bar graph. Error bars reflect standard error across different lesion locations within each lesion syndrome. (B) and (C) Lesions (white outlines) associated with confabulation (showing 4 of 25 cases) and criminal behavior (showing 4 of 17 cases) showed the strongest intersections with positive nodes of our religious fundamentalism network, similar to lesions associated with high religious fundamentalism. (D) Lesion locations associated with post-stroke pain (showing 4 of 23 cases) showed the strongest intersection with negative nodes of our spirituality circuit, similar to lesion locations low religious fundamentalism.

In addition, we observed strong spatial similarities between the fundamentalism network and brain lesion connectivity associated with criminal behavior (spatial correlations: t_16_ = 5.5, *p* < 10^-4^, 95% CI, *r =* 0.28 to 0.63; Figure 5). This is consistent with previously observed associations between high religious fundamentalism and increased hostility, aggression, and violence against outgroups^8,18,44^.

Finally, strong negative spatial correlations were observed between our religious fundamentalism network and lesion connectivity associated with central post-stroke pain (spatial correlations: t_22_ = -5.2, *p* < 10^-4^, 95% CI, *r =* -0.4462 to -0.1926; Figure 5). This inverse association is consistent with prior work showing that religious stimuli may engage pain-inhibiting brain processes and induce analgesic effects^45^. These data suggest that neurobiological factors in healthy individuals predict a style of cognitive and emotional processing that often result in religious fundamentalism.

## Discussion

Brain lesions associated with religious fundamentalism occur in different brain regions but can be considered components of a single, connected brain network. This religious fundamentalism network was consistent and cross-validated in two large, etiologically diverse neurological patient samples collected from independent laboratories. It incorporated regions in the right superior orbital frontal, right middle frontal, right inferior parietal, right inferior temporal cortices, and left cerebellum. Our lesion network mapping findings were reproducible across independent datasets, and networks derived from one dataset could be used to predict religious fundamentalism scores in the opposite dataset. The cross-dataset replication and prediction depended on network mapping, as it was not present using lesion locations alone. Moreover, lesions to regions in the fundamentalism network were positively associated with confabulation and criminal behavior^46^ and negatively associated with pain.

Given the etiological differences between our patient samples in the two datasets, our robust replication of the fundamentalism brain network is remarkable. One sample included Vietnam war veterans with penetrating traumatic brain injury^47^, and the other sample was neurological patients drawn from the Iowa Neurological Patient Registry^48^ who sustained focal brain injury mainly from stroke or surgical resection for treatment of a meningioma or seizure control. Traumatic brain injury, slow-growth meningioma resections, or stroke injury can all lead to distinct sequela of symptoms following damage to the same region^49^. Our reproduction and cross-validation across these etiologically diverse datasets increase our confidence that lesions within a well-defined network change the probability of religious fundamentalism.

Moreover, the right cerebral cortex lateralization of our fundamentalism network is consistent with split-brain and other unilateral lesion evidence regarding reasoning. Whereas the left hemisphere draws inferences, hypotheses, and rationalizations from the evidence available to it, the right hemisphere stops the perseveration of incorrect ideas^50–52^. Right hemisphere damage can produce confabulatory or delusional beliefs^53–55^ with impairments in error monitoring or conflict detection. In the religious domain, right hemisphere damage may result in reduced recognition of belief conflict, less religious disbelief, and the persistence of more extreme religious beliefs.

It is important not to overinterpret our brain network results. Many factors contribute to fundamentalism, including affective, cognitive, experiential, genetic, familial, institutional, developmental, and cultural variables. Given these multiple factors, some patients with lesions in our religious fundamentalism network may not display high fundamentalism. Although brain lesions to this network may increase the likelihood of religious fundamentalism^6^, they should not be interpreted as an inevitable or sole cause of fundamentalism. Moreover, the reverse inference that individuals high in religious fundamentalism have brain lesions is unsubstantiated and unwarranted. Similarly, our lesion results do not imply that people with strong religious beliefs confabulate or that individuals high in religious fundamentalism commit crimes. Rather, our data may help us understand the style of cognitive or emotional processing that increase or decrease the probability of holding fundamentalism attitudes.

A limitation in the current work is that patients in our study are predominantly older Caucasian individuals from Christian backgrounds. This may limit generalizability to populations with more variabilities in age, gender, ethnicity, culture, and religious tradition. Future studies with diverse patient samples are warranted to validate the generalizability of these conclusions.

In sum, brain lesions associated with greater religious fundamentalism are characterized by a unique pattern of brain connectivity: a religious fundamentalism network. Specifically, lesion connectivity to the right orbitofrontal cortex, dorsolateral prefrontal cortex and inferior parietal lobes corresponds to high fundamentalism attitudes. Our lateralized result is consistent with well-established hemispheric specializations in reasoning. This network spatially resembles brain lesion networks associated with pathological confabulation and criminal behavior. Overall, these results unify previously published reports on the neural substrates for religious fundamentalism and may prove useful for understanding neurobiological factors that influence the development of fundamentalism and its associated cognitive and emotional profile.

## Materials and Methods

### Subjects

One hundred and ninety patients with brain damage from the W.F. Caveness Vietnam Head Injury Study Registry (VHIS) and the Iowa Neurological Patient Registry comprised the two datasets of our study. Dataset 1 had 106 patients with penetrating traumatic brain injury who were male combat veterans from VHIS during phase 4 (40 years after injury) conducted at the National Institute of Neurological Disorders and Stroke (NINDS) in Bethesda, MD^47^. Patients in dataset 1 self-identified religious affiliations included Protestant (39%), Roman Catholic (16%), other affiliation (12%), and 33% did not identify with a religious affiliation^7^. Dataset 2 had 84 patients from a demographically homogenous region (mainly rural Iowa) who experienced brain lesions from diverse etiologies: stroke (51%), tumor or seizure control surgical brain resection (37%), traumatic brain injury (8%), and 4% had brain damage from a viral infection or a genetic condition. Patients in dataset 2 self-identified religious affiliations included Protestant (44%), Roman Catholic (25%), other affiliation (18%), and 13% did not identify with a religious affiliation. All participants gave written informed content prior to data collection.

### Religious fundamentalism assessment

Religious fundamentalism was measured using the Religious Fundamentalism Scale^3,4^ (RFS). The RFS is a widely-used, well-validated, and standardized psychometric instrument assessing religious fundamentalism. Patients recorded responses to RFS statements on a Likert scale (*Strongly agree* to *Strongly disagree*). The RFS defines fundamentalism with 4 dimensions: 1) there is one set of religious teachings that contain the fundamental, inerrant truth about humanity and deity; 2) this truth is opposed to evil which must be actively fought; 3) this truth must be followed today according to the fundamental practices of the past; and 4) those who follow these fundamental teachings have a special relationship with the deity. Patients in dataset 1 were administered a randomized and balanced 10-item version of the 1992 RFS^3,7^ while patients in dataset 2 were administered the balanced 12-item 2004 version of the RFS^3,6^. The balanced items of the scale rule out the possibility that liberal responding, per se, produces a high or low fundamentalism score.

### Neuroanatomical analyses

Neuroimaging data in all patients was obtained in the chronic epoch, 3 or more months following lesion onset. Brain lesion neuroanatomical identification in dataset 1 patients was based on computerized tomography (CT) scans for each patient^7^. The Analysis of Brain Lesion (ABLe) software version 2.8b^56^ implemented in MEDx version 3.44 (Medical Numerics, Germantown, MD) was used to calculate lesion location and volume loss. Each CT scan was then spatially normalized to a CT template brain image in Montreal Neurological Institute (MNI-152) template space (resolution, 1 mm^3^) by applying an automated image registration algorithm^57^ for registration accuracy. Brain lesion neuroanatomical identification in dataset 2 was based on either on CT or magnetic resonance imaging (MRI) scans for each patient. Each patient’s lesion was reconstructed in three dimensions using Brainvox, and the lesion contour was manually warped into a template brain creating a mask using the MAP-3 method^58^. Lesion masks were warped and resampled from the template brain local standard space to MNI-152 template space using a symmetric normalization algorithm in ANTS^59^. All lesions in both datasets were aligned to the same MNI-152 template space.

### Lesion network mapping

To maximize the statistical power in our analysis, we combined both independent datasets (*N*1 = 106; *N*2 = 84) and performed voxelwise statistical tests for lesion network associations with RFS scores (*N*_combined_ = 190). We used lesion network mapping and previously validated methods to derive a brain network for religious fundamentalism in a data-driven fashion^2^. First, resting-state functional connectivity between each lesion and the rest of the brain was computed using a publicly available normative connectome Dataset from 1000 healthy right-handed subjects (42.7% male subjects, ages 18–35 years, mean age 21.3 years). This connectome dataset was processed in accordance with the lesion network mapping procedure^60^ which results in a map of brain regions functionally connected to each lesion location referred to as a lesion network. In a voxelwise fashion for both independent datasets, a group-level map of brain associations with religious fundamentalism was derived. This was done using voxelwise permutation analysis of linear models (PALM) with RFS scores as a psychometric covariable (Figure 2). We performed a family-wise error (FWE) multiple comparison correction on our results and tested for voxels that survived a conservative correction threshold of FWE p < 0.05. Coordinates for peak values in clusters surviving FWE correction were identified using MRIcroGL and the Automated Anatomical Labeling (AAL) atlas.

### Cross-validations analyses

Cross-validation testing was performed in two ways. First, spatial correlations were calculated between each individual patients’ brain lesion network maps compared with the group-level fundamentalism network map from the opposing dataset. Specifically, the group-level fundamentalism map derived from dataset 2 was spatially correlated with each individual patient’s lesion network map from dataset 1, and the group-level fundamentalism map derived from dataset 1 was spatially correlated with each individual patient’s lesion network map from dataset 2. This quantified the similarity of each patient’s lesion network to an independent group-level fundamentalism network derived from the opposing dataset. Each of these patient-to-group network similarity scores were then correlated with patients’ own RFS scores to test whether the similarity of an individual patients’ lesion network map to an independently derived fundamentalism network corresponded to patients’ psychometric scores for religious fundamentalism. Second, we calculated a network damage score for each patients’ brain lesion location relative to the opposite datasets’ group-level fundamentalism network. This was done by superimposing each patient’s brain lesion image onto the fundamentalism network map from the opposite dataset, then calculating the arithmetic sum of the statistical values for each voxel in the group-level fundamentalism map that was circumscribed by the patient’s lesion image^2^. These network damage scores were then correlated with the patients’ RFS scores to test whether focal damage to a fundamentalism network map corresponded to patients’ psychometric scores for religious fundamentalism.

### Religious fundamentalism network and lesions associated with behavioral, neurological, and psychiatric conditions analyses

We compared the religious fundamentalism brain network to independent sets of previously published brain lesions and lesion networks associated with behavioral, neurological and psychiatric conditions from a library of 899 lesions spanning 21 conditions. We used two methods of comparison in these analyses: First, we assessed the extent of the intersection between individual condition-associated lesions and the unthresholded fundamentalism network to compute a “network damage score”^2^: individual condition-associated lesions were superimposed on the combined dataset religious fundamentalism network, and the average of the t-value for voxels circumscribed within each lesion was calculated. A one-sample t-test was then performed for network damage score values associated with brain lesion traces within a set of lesions sharing a common symptom. These t-test results were then corrected for multiple comparisons across the 21 sets of condition-associated lesions. Second, we compared the spatial topography of the individual lesion network maps from each condition to the unthresholded religious fundamentalism network. Degree of spatial similarity to the fundamentalism network with condition-associated lesion networks was then calculated by performing a Pearson correlation between vectorized maps of our religious fundamentalism network and each of the lesion network maps for the 899 condition-associated lesions in our brain lesion library. A one-sample t-test was then performed for spatial similarity values associated with lesion networks within a set of lesions sharing a common condition. These t-test results were then corrected for multiple comparisons across the 21 sets of condition-associated lesions.

## Acknowledgements

While most scientific papers include limitations, the authors would like to add a note of humility. In our personal beliefs, we span a broad continuum from adherents of religious faiths through agnosticism to atheism. We approach this weighty subject matter as earnest seekers and hope that you receive our results in the spirit of open-minded empirical inquiry driven by scientific curiosity and without prejudice or malice to any group or faith.

---

**Supplementary Figure 1:**
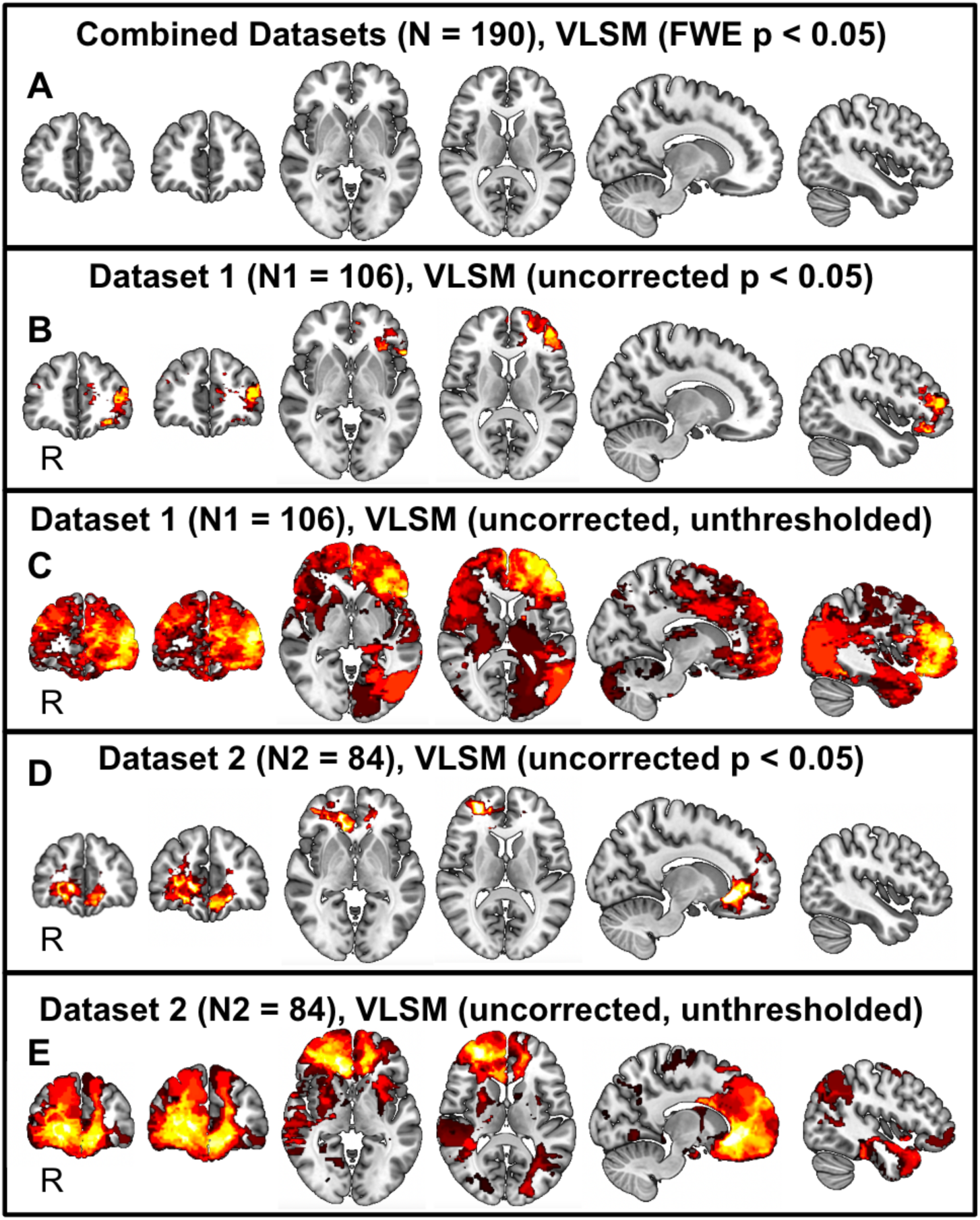
Voxelwise lesion-symptom mapping (VLSM) for religious fundamentalism. (A) No voxels survive Family-Wise Error (FWE) multiple comparison correction in a combined dataset analysis (*N* = 190) testing for associations between fundamentalism and lesion location. (B & C) Statistically uncorrected peak associations between religious fundamentalism and brain lesion locations determined by VLSM in Dataset 1 (*N*1 = 106) localized to left lateral prefrontal cortex. (D & E) Statistically uncorrected peak associations between religious fundamentalism and brain lesion locations determined by VLSM in Dataset 2 (*N*2 = 84) localized to the right medial prefrontal cortex. Attempted cross-validation of VLSM findings between Datasets 1 and 2 failed to achieve statistical significance for either thresholded or unthresholded VLSM results.

## SUPPLEMENTARY METHODS

### Voxel lesion-symptom mapping (VLSM) in Datasets 1 and 2

To test whether religious fundamentalism could be explained by a data-driven analysis of lesion locations rather than lesion networks, voxel lesion-symptom mapping (VLSM) was performed. To parallel with our lesion network mapping methods, VLSM was performed by combining Datasets (*N*_combined_ = 190) to maximize statistical power, and on independent Datasets (*N*1 = 106; *N*2 = 84) to test for replication using religious fundamentalism scores as covariates with lesion locations. Univariate voxel-based lesion-symptom mapping (VLSM) in PALM and multivariate VLSM^50^ in the SVR-LSM toolbox (https://github.com/atdemarco/svrlsmgui) were both applied. Univariate VLSM applied a general linear regression model and permutation test, controlling for covariates. We limited our analysis to voxels occurring in at least 5% of lesions, assessed with Freedman-Lane permutations (the default setting of 1000 permutations was used) while controlling for lesion volume as a covariate and correcting for multiple testing. These parameters were chosen based on published best-practice recommendations^61, 62^. A two-tailed family wise error corrected p-value < 0.05 was considered significant. SVR-LSM performs multivariate voxel- and cluster-based LSM with a machine learning regression, termed the support vector regression (SVR), whereas univariate VLSM considers neighboring voxels as independent. Multivariate SVR-VLSM simultaneously considers many voxels at once when determining whether damaged brain regions contribute to behavioral measures^63, 64^. These multivariate LSM approaches can identify complex dependencies that traditional univariate VLSM approaches cannot. In line with the univariate VLSM analysis, we limited our analysis for multivariate VLSM to voxels occurring in at least 5% of lesions, assessed with permutations (default setting of 1,000 permutations was used), while controlling for lesion volume using the standard “direct total lesion volume control” (dTLVC) approach and correcting for multiple testing. A two-tailed family wise error corrected p-value < 0.005 was considered significant. These parameters were chosen based on published best-practice recommendations^63, 64, 65^.

Additionally, we explored univariate and multivariate VLSM results using liberal statistical cutoffs and assessed their alignment with our findings from lesion-network mapping by overlapping statistically significant VLSM voxels with lesion network mapping results. Restricted maps were derived by applying a voxelwise threshold of p < 0.05 to regions with a minimum of 5% lesion overlap. A more liberal unrestricted and unthresholded map was also derived for each Dataset, also using religious fundamentalism scores as covariates with lesion locations.

Both the univariate and the multivariate VLSM analyses were applied to Datasets 1 and 2 separately (*N*1 = 106; *N*2 = 84), and to combined Datasets to boost statistical power through increased sample size (*N*_combined_ = 190).

### Attempted cross-validation of voxel lesion-symptom mapping (VLSM) results

Cross-validation testing of VLSM results was performed by, first, superimposing brain lesion locations from Dataset 1 (*N*1 = 106) onto both the restricted and the unrestricted maps derived from Dataset 2 (*N*2 = 84). Second, a “network damage score” was calculated for each individual lesions: the “network damage score” is defined as the arithmetic sum of the statistical values for each voxel in the VLSM reference map that is circumscribed by a given lesion. These “network damage scores” for each patient were then correlated with the patients’ religious fundamentalism scores. Any significant correlations between the “network damage scores” and the religious fundamentalism scores would be considered evidence of successful cross-validation between Datasets.

## References

1 Kapogiannis, D., Barbey, A. K., Su, M., Zamboni, G., Krueger, F., & Grafman, J. (2009). Cognitive and neural foundations of religious belief. Proceedings of the National Academy of Sciences, 106(12), 4876–4881.

2 Ferguson, M. A. et al. A neural circuit for spirituality and religiosity derived from patients with brain lesions. Biological Psychiatry 91, 380–388 (2022).

3 Altemeyer, B. & Hunsberger, B. Authoritarianism, religious fundamentalism, quest, and prejudice. The International Journal for the Psychology of Religion 2, 113–133 (1992).

4 Altemeyer, B. & Hunsberger, B. A revised religious fundamentalism scale: The short and sweet of it. The International Journal for the Psychology of Religion 14, 47–54 (2004).

5 Altemeyer, B. & Hunsberger, B. in Handbook of the Psychology of Religion and Spirituality (eds R.F. Paloutzian & C.L. Park) 378–393 (Guliford Press, 2005).

6 Asp, E. W., Ramchandran, K. & Tranel, D. Authoritarianism, religious fundamentalism, and the human prefrontal cortex. Neuropsychology 26, 414–421 (2012).

7 Zhong, W., Cristofori, I., Bulbulia, J., Krueger, F. & Grafman, J. Biological and cognitive underpinnings of religious fundamentalism. Neuropsychologia 100, 18–25 (2017).

8 Altemeyer, B. The Authoritarian Specter. (Harvard University Press, 1996).

9 Zmigrod, L., Rentfrow, P. J., Zmigrod, S. & Robbins, T. W. Cognitive flexibility and religious disbelief. Psychological Research 83, 1749–1759 (2019).

10 Weeks, M. & Geisler, S. Revisiting cognitive complexity of religious topics: Multiple complexity, religious fundamentalism, and a quest orientation. Psychology of Religion and Spirituality 11, 433–441 (2019).

11 Hunsberger, B., Alisat, S., Pancer, S. M. & Pratt, M. Religious fundamentalism and religious doubts: Content, connections, and complexity of thinking. The International Journal for the Psychology of Religion 6, 201–220 (1996).

12 Asp, E. W. & Tranel, D. in Principles of Frontal Lobe Function (eds D.T. Stuss & R.T. Knight) 383–416 (Oxford University Press, 2013).

13 Bronstein, M. V., Pennycook, G., Bear, A., Rand, D. G. & Connon, T. D. Belief in fake news is associated with delusionality, dogmatism, religious fundamentalism, and reduced analytic thinking. Journal of Applied Research in Memory and Cognition 8, 108–117 (2019).

14 Sobol, M., Zajenkowski, M. & Jankowski, K. S. Religious fundamentalism, delusions, and conspiracy beliefs related to the COVID-19 pandemic. International Journal of Environmental Research and Public Health 19, 1–6 (2022).

15 Merkley, E. & Loewen, P. J. Anti-intellectualism and the mass public’s response to the COVID-19 pandemic. Nature Human Behaviour 5, 706–715 (2021).

16 Oliver, J. E. & Rahn, W. M. Rise of the Trumpenvolk populism in the 2016 election. The Annals of the American Academy of Political and Social Science 667, 189–206 (2016).

17 Saroglou, V. et al. Fundamentalism as dogmatic belief, moral rigorism, and strong groupness across cultures: Dimensionality, underlying components, and related interreligious prejudice. Psychology of Religion and Spirituality, 1–14 (2020).

18 Koopmans, R. Religious fundamentalism and hostility against out-groups: A comparison of Muslims and Christians in Western Europe. Journal of Ethnic and Migration Studies 41, 33–57 (2015).

19 Jackson, L. M. & Hunsberger, B. An intergroup perspective on religion and prejudice. Journal for the Scientific Study of Religion 38, 509–523 (1999).

20 Lueke, N. A., Lueke, A. K., Aghababaei, N., Ferguson, M. A. & Bushman, B. J. Fundamentalism and intrinsic religiosity as factors in well-being and social connectedness: An Iranian study. Psychology of Religion and Spirituality, 1–9 (2021).

21 Beer, J. M., Arnold, R. D. & Loehlin, J. C. Genetic and environmental influences on MMPI factor scales: Joint model fitting to twin and adoption data. Journal of Personality and Social Psychology 74, 818–827 (1998).

22 Coreno, T. Fundamentalism as a class culture. Sociology of Religion 63, 335–360 (2002).

23 Wilson, J. & Sandomirsky, S. Religious affiliation and the family. Sociological Forum 6, 289–309 (1991).

24 Devinsky, O. & Lai, G. Spirituality and religion in epilepsy. Epilepsy & Behavior 12, 636–643 (2008).

25 Griffiths, R. R., Richards, W. A., McCann, U. & Jesse, R. Psilocybin can occasion mystical-type experiences having substantial and sustained personal meaning and spiritual significance. Psychopharmacology 187, 268–283 (2006).

26 Azari, N. P. et al. Neural correlates of religious experience. European Journal of Neuroscience 13, 1649–1652 (2001).

27 Beauregard, M. & Paquette, V. EEG activity in Carmelite nuns during a mystical experience. Neuroscience Letters 444, 1–4 (2008).

28 Moll, J., Krueger, F., Zahn, R., Pardini, M., de Oliveira-Souza, R., & Grafman, J. (2006). Human fronto–mesolimbic networks guide decisions about charitable donation. Proceedings of the National Academy of Sciences, 103(42), 15623–15628.

29 Koenig, L. B., McGue, M., Krueger, R. F. & Bouchard, T. J. Genetic and environmental influences on religiousness: Findings for retrospective and current religiousness ratings. Journal of Personality 73, 471–488 (2005).

30 Bouchard, T. J. et al. in Behavior genetic principles: Perspectives in Development, Personality, and Psychopathology (ed L.F. DeLilla) 89–104 (American Psychological Association, 2004).

31 Ludeke, S., Johnson, W. & Bouchard, T. J. "Obedience to traditional authority:" A heritable factor underlying authoritarianism, conservatism and religiousness. Personality and Individual Differences 55, 375–380 (2013).

32 Pennycook, G., Tranel, D., Warner, K. & Asp, E. W. in Neurology and Religion (eds A. Coles & J. Collicutt) 115–129 (Cambridge University Press, 2019).

33 Siddiqi, S. H., Kording, K. P., Parvizi, J. & Fox, M. D. Causal mapping of human brain function. Nature Reviews Neuroscience 23, 361–375 (2022).

34 Vaidya, A. R., Pujara, M. S., Petrides, M., Murray, E. A. & Fellows, L. K. Lesion studies in contemporary neuroscience. Trends in Cognitive Sciences 23, 655–673 (2019).

35 Urgesi, C., Aglioti, S. M., Skrap, M. & Fabbro, F. The spiritual brain: Selective cortical lesions modulate human self-transcendence. Neuron 65, 309–319 (2010).

36 Petrides, M. & Pandya, D. N. in Principles of Frontal Lobe Function (eds D.T. Stuss & R.T. Knight) 31-50 (Oxford University Press, 2002).

37 Uddin, L. Q., Yeo, B. T. & Spreng, R. N. Towards a universal taxonomy of macro-scale functional human brain networks. Brain Topography 32, 926–942 (2019).

38 Boes, A. D. et al. Network localization of neurological symptoms from focal brain lesions. Brain 138, 3061–3075 (2015).

39 Palesi, F. et al. Contralateral cortico-ponto-cerebellar pathways reconstruction in humans in vivo: Implications for reciprocal cerebro-cerebellar structural connectivity in motor and non-motor areas. Scientific Reports 7, 12841 (2017).

40 Bateman, J. R. et al. Network localization of spontaneous confabulation. The Journal of Neuropsychiatry and Clinical Neurosciences, 1–8 (2023).

41 Moscovitch, M. in Varieties of Memory and Consciousness: Essays in Honour of Endel Tulving (eds H.L. Roediger & F. Craik) 133–160 (Erlbaum, 1989).

42 Baddeley, A. & Wilson, B. in Autobiographical Memory (ed D. Rubin) 225–252 (Cambridge University Press, 1986).

43 Ciaramelli, E., Ghetti, S., Frattarelli, M. & Làdavas, E. When true memory availability promotes false memory: Evidence from confabulating patients. Neuropsychologia 44, 1866–1877 (2006).

44 Juergensmeyer, M. Terror in the mind of God: The global rise of religious violence. 4th edn, (University of California Press, 2017).

45 Wiech, K. et al. An fMRI study measuring analgesia enhanced by religion as a belief system. Pain 139, 467–476 (2008).

46 Darby, R. R., Horn, A., Cushman, F., & Fox, M. D. (2018). Lesion network localization of criminal behavior. Proceedings of the National Academy of Sciences, 115(3), 601–606.

47 Raymont, V., Salazar, A. M., Krueger, F. & Grafman, J. "Studying Injured Minds" -- The Vietnam Head Injury Study and 40 years of brain injury research. Frontiers in Neurology 2, 1–13 (2011).

48 Tranel, D. in Textbook of Clinical Neuropsychology (eds J.E. Morgan & J.H. Ricker) 27–39 (Taylor and Francis, 2007).

49 Damasio, H. in Handbook of Neuropsychology Vol. 1 (eds F. Boller, J. Grafman, & G. Rizzolatti) 77-102 (Elsevier, 2000).

50 Turner, B. O., Marinsek, N., Ryhal, E. & Miller, M. B. Hemispheric lateralization in reasoning. Annals of the New York Academy of Sciences 1359, 47–64 (2015).

51 Gazzaniga, M. S. in Self and consciousness 88-102 (Psychology Press, 2014).

52 Wolford, G., Miller, M. B. & Gazzaniga, M. S. The left hemisphere’s role in hypothesis formation. The Journal of Neuroscience 20, 1–4 (2000).

53 Marcel, A. J., Tegner, R. & Nimmo-Smith, I. Anosognosia for plegia: specificity, extension, partiality, and disunity of bodily unawareness. Cortex 40, 19–40 (2004).

54 Feinberg, T. E., Roane, D. M., Kwan, P. C., Schindler, R. J. & Haber, L. D. Anosognosia and visuoverbal confabulation. Archives of Neurology 51, 468–473 (1994).

55 Vocat, R., Staub, F., Stroppini, T. & Vuilleumier, P. Anosognosia for hemiplegia: A clinical-anatomical prospective study. Brain 133, 3578–3597 (2010).

56 Makale, M. et al. Quantification of brain lesions using interactive automated software. Behavior Research Methods, Instruments, & Computers 34, 6–18 (2002).

57 Woods, R. P., Grafton, S. T., Holmes, C. J., Cherry, S. R. & Mazziotta, J. C. Automated image registration: I. General methods and intrasubject, intramodality validation. Journal of Computer Assisted Tomography 22, 139–152 (1998).

58 Frank, R. J., Damasio, H. & Grabowski, T. J. Brainvox: An interactive, multimodal visualization and analysis system for neuroanatomical imaging. Neuroimage 5, 13–30 (1997).

59 Avants, B. & Gee, J. C. Geodesic estimation for large deformation anatomical shape averaging and interpolation. Mathematics in Brain Imaging 23, S139–S150 (2004).

60 Fox, M. D. Mapping symptoms to brain networks with the human connectome. The New England Journal of Medicine 379, 2237–2245 (2018).

## SUPPLEMENTARY REFERENCES

61 Sperber, C. & Karnath, H.O. Impact of correction factors in human brain lesion-behavior inference. Human Brain Mapping 38(3), 1692–1701 (2017).

62 Karnath, H.O., Sperber, C. & Rorden, C. Mapping human brain lesions and their functional consequences. Neuroimage 165, 180–189 (2018).

63 Zhang Y, Kimberg DY, Coslett HB, Schwartz MF, & Wang Z. Multivariate lesion-symptom mapping using support vector regression. Human Brain Mapping 35(12), 5861–5876 (2014).

64 DeMarco, A.T. & Turkeltaub, P.E. A multivariate lesion symptom mapping toolbox and examination of lesion-volume biases and correction methods in lesion-symptom mapping 39(11), 4169-4182). Hoboken, USA: John Wiley & Sons, Inc (2018).

65 Mah Y-H, Husain M, Rees G, & Nachev P. Human brain lesion-deficit inference remapped. Brain 137, 2522–2531 (2014).

